# Exchange catalysis during anaerobic methanotrophy revealed by ^12^CH_2_D_2_ & ^13^CH_3_D in methane

**DOI:** 10.1101/377531

**Authors:** Jeanine L. Ash, Matthias Egger, Tina Treude, Issaku Kohl, Barry Cragg, R. John Parkes, Caroline P. Slomp, Barbara Sherwood Lollar, Edward D. Young

## Abstract

The anaerobic oxidation of methane (AOM) is a crucial component of the methane cycle, but its enzymatic versatility under environmental conditions remains poorly understood. We use sediment samples collected during IODP Expedition 347 to the Baltic Sea to show that relative abundances of ^12^CH_2_D_2_ and ^13^CH_3_D molecules in methane gas trace the reversibility of methyl-coenzyme M reductase during AOM by driving methane towards internal, thermodynamic isotopic equilibrium. These data suggest that ^12^CH_2_D_2_ and ^13^CH_3_D together can identify the influence of methanotrophy in environments where conventional bulk isotope ratios are ambiguous, and these findings may lead to new insights regarding the global significance of enzymatic back-flux in the methane cycle.

---

Quantifying the subsurface carbon cycle by linking biogeochemistry to deep biosphere metabolisms remains a long-standing challenge compounded by slow rates of reaction (WEBSTER *et al.*, 2015; WAITE *et al.*, 2017). Even in well-characterized environments, determining how methanogens and anaerobic methanotrophs influence organic matter degradation can be challenging (REGNIER *et al.*, 2011; BEULIG *et al.*, 2018a). These challenges are due in part to how methane produced by methanogens inherits characteristics from its substrates (both a range of organic and inorganic carbon) (CONRAD, 2005), and they are further complicated when methane is itself the substrate during anaerobic methane oxidation (AOM) (WHITICAR, 1999). Our understanding of how anaerobic methanotrophs respond to varying environmental conditions (HOLLER *et al.*, 2009; HOLLER *et al.*, 2011; YOSHINAGA *et al.*, 2014) and how they might contribute to “cryptic cycles” (the rapid production and consumption of reactive intermediates) in the marine subsurface is constantly evolving (EGGER *et al.*, 2018).

AOM is thought to be carried out by a consortium of methane-oxidizing archaea and sulphate-reducing bacteria performing the net reaction 
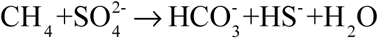
 (BOETIUS *et al.*, 2000). Evidence suggests that these processes can be decoupled (MILUCKA *et al.*, 2012; SCHELLER *et al.*, 2016), and metal oxides may also act as terminal electron acceptors (BEAL *et al.*, 2009; CAI *et al.*, 2018). In all cases, the first step of AOM involves C-H bond activation by a modified methyl-coenzyme M reductase (Mcr), the terminal enzyme in methanogenesis (MILUCKA *et al.*, 2012; SCHELLER *et al.*, 2016). Anaerobic methanotrophs use enzymes homologous to those used by methanogens, prompting questions regarding the directionality of these metabolisms (LLOYD *et al.*, 2011; BEULIG *et al.*, 2018b).

Coupling exploration of methane enzymatic chemistry with isotope geochemistry can provide powerful insights. Here, we interrogate subsurface methane metabolisms by precisely determining concentrations of molecules containing two heavy isotopes, referred to as “clumped” isotopologues (EILER, 2007). The relative proportions of ^13^CH_3_D and ^12^CH_2_D_2_ (reported as Δ^13^CH_3_D and Δ^12^CH_2_D_2_ values relative to a random distribution of isotopes among all CH_4_ isotopologues; see Supplementary Information) have known temperature-sensitive equilibrium concentrations and thus are sensitive indicators of reversibility in reactions (PIASECKI *et al.*, 2016). For instance, axenic cultures of methanogens produce methane with depletions in clumped isotopes relative to the random distribution that reflect kinetic processes (STOLPER *et al.*, 2015; WANG *et al.*, 2015; DOUGLAS *et al.*, 2016; YOUNG *et al.*, 2017; GRUEN *et al.*, 2018). In some cases, the clumped isotope composition of environmental microbial methane is consistent with the axenic cultures, but in others it displays a clumped isotope composition reflective of equilibration within the environment of formation (WANG *et al.*, 2015; YOUNG *et al.*, 2017). This span in clumped-isotope distributions suggests that enzymatic reactions associated with methanogenesis and methanotrophy are capable of ranging from reversible to kinetic. Previous work using one rare isotopologue (Δ^13^CH_3_D) suggests that that rate of methanogenesis controls the degree of molecular isotopic equilibrium (STOLPER *et al.*, 2015; WANG *et al.*, 2015). However, the inability to produce equilibrated methane from axenic cultures has led to speculation that AOM may be responsible for equilibrated environmental methane (STOLPER *et al.*, 2015; WANG *et al.*, 2016; YOUNG *et al.*, 2017; GRUEN *et al.*, 2018). The use of a second rare isotopologue, ^12^CH_2_D_2_, aids in interpreting the causes of intra-methane disequilibrium and suggests that microbial methanogenesis does not lead to isotopologue equilibrium among methane molecules.

Here, we examine microbial methane from the Bornholm Basin (Baltic Sea; see Supplementary Information) to test the hypothesis that enzymatic-back reaction during AOM drives methane to thermodynamic equilibrium. We compare methane clumped isotope composition to geochemical profiles (ANDRÉN *et al.*, 2015; EGGER *et al.*, 2017) and present-day rates of methanogenesis and methanotrophy calculated with a multicomponent diagenetic model (DIJKSTRA *et al.*, 2018) to determine parameters controlling the isotopic composition of methane (Fig 1A-D). Methane δ^13^C and δD, δ^13^C_DIC_, δ^13^C_TOC_ and δD_H2O_ vary non-monotonically over 30 meters below seafloor (Fig. 1H-G, Fig. S1) while Δ^13^CH_3_D and Δ^12^CH_2_D_2_ values increase with depth (Fig. 1E-F).

**Figure 1:**
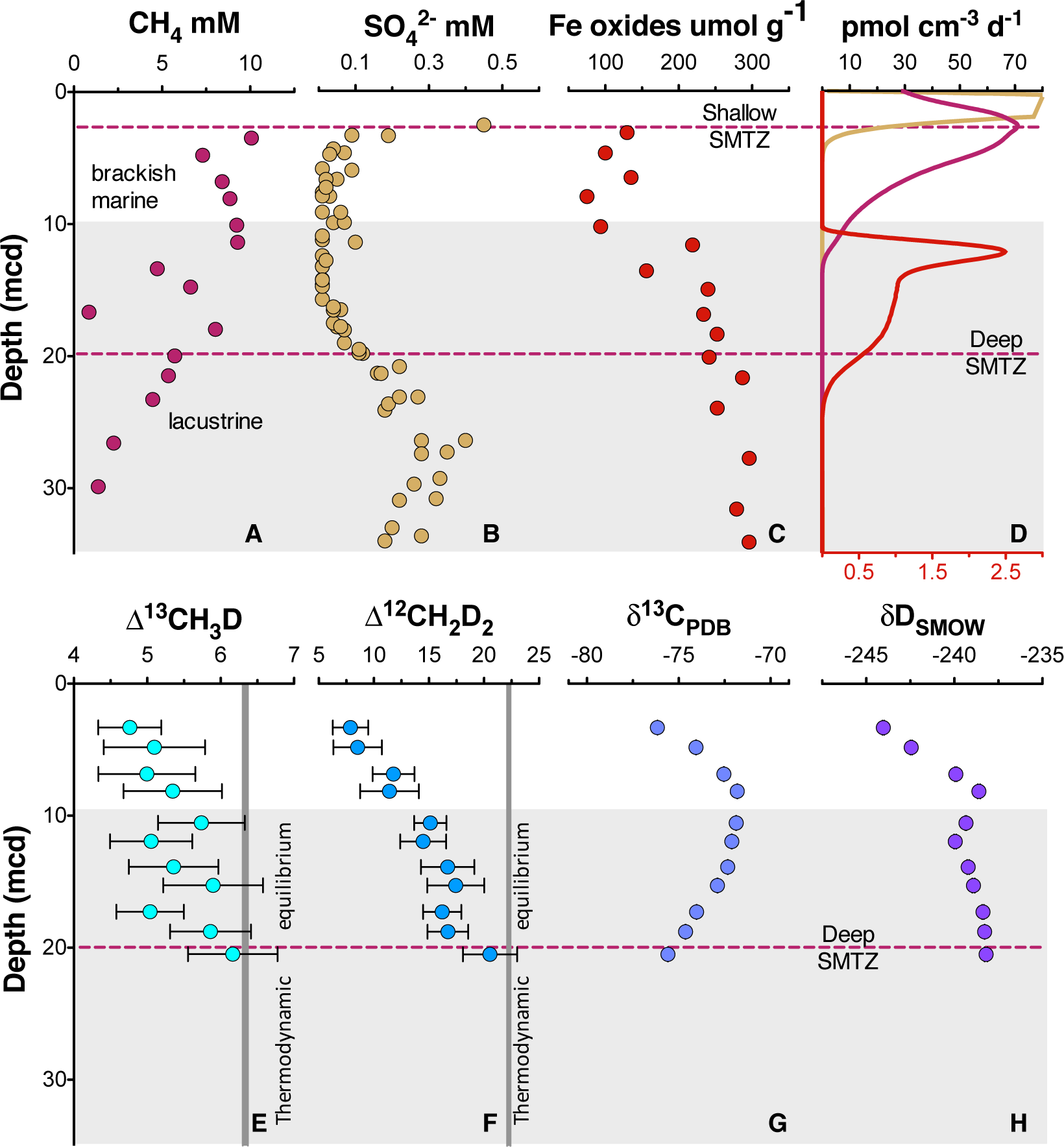
Geochemical profiles (±2σ) from Bornholm Basin from 3-35 MCD. Horizontal grey bar denotes sediments deposited during lacustrine conditions overlain with sediments deposited during brackish marine conditions. Samples were not recovered above the shallow SMTZ. Pink dashed lines denote a shallow (3.3 MCD) and deep (19.8 MCD) SMTZ; only the shallow SMTZ is shown in the bottom panel for clarity. Dark grey vertical bars in **E** and **F** denote equilibrium isotopologue compositions for the subsurface temperature of 7.8 ± 0.6°C. Modelled CH_4_ production, Fe-AOM and SO_4_-AOM rates are shown in panel **D** (DIJKSTRA *et al.*, 2018). CH_4_ production (purple) and SO_4_-AOM (gold) rates correspond with the upper x-axis and Fe-AOM rates (red) correspond with the lower x-axis. SO_4_-AOM rates above the upper SMTZ are calculated using bottom-water SO_4_ concentrations of 15.0 mmol. Additional porewater data shown in Supplemental Information.

Comparisons of the bulk isotopic composition of methane to carbon and hydrogen reservoirs at this location are complicated by the change from limnic to marine deposition (EGGER *et al.*, 2017). For instance, the hydrogen isotopes in methane are seemingly not equilibrated with porewater that changes by >100‰ in this interval (Fig. S1), obscuring the history of this reservoir. Instead, we will focus on the intra-molecular isotopic equilibrium of the methane itself using clumped isotopes, thus sidestepping thecomplications in bulk isotope ratios. We introduce 
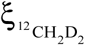
, a parameter that quantifies the degree of thermodynamic disequilibrium for a methane system. 
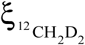
 is analogous to the symbol for the reaction progress variable (
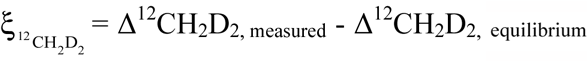
, see Supplementary Information). Values for 
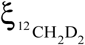
 in upper sediments as low as −7.5‰ are significantly different (>6σ) from 0‰ (*i.e.* thermodynamic equilibrium) which is approached with depth (Fig. 2). In order to determine how methanogenesis and methanotrophy influence this transition, we consider evidence for and against reversibility in each metabolism.

**Figure 2:**
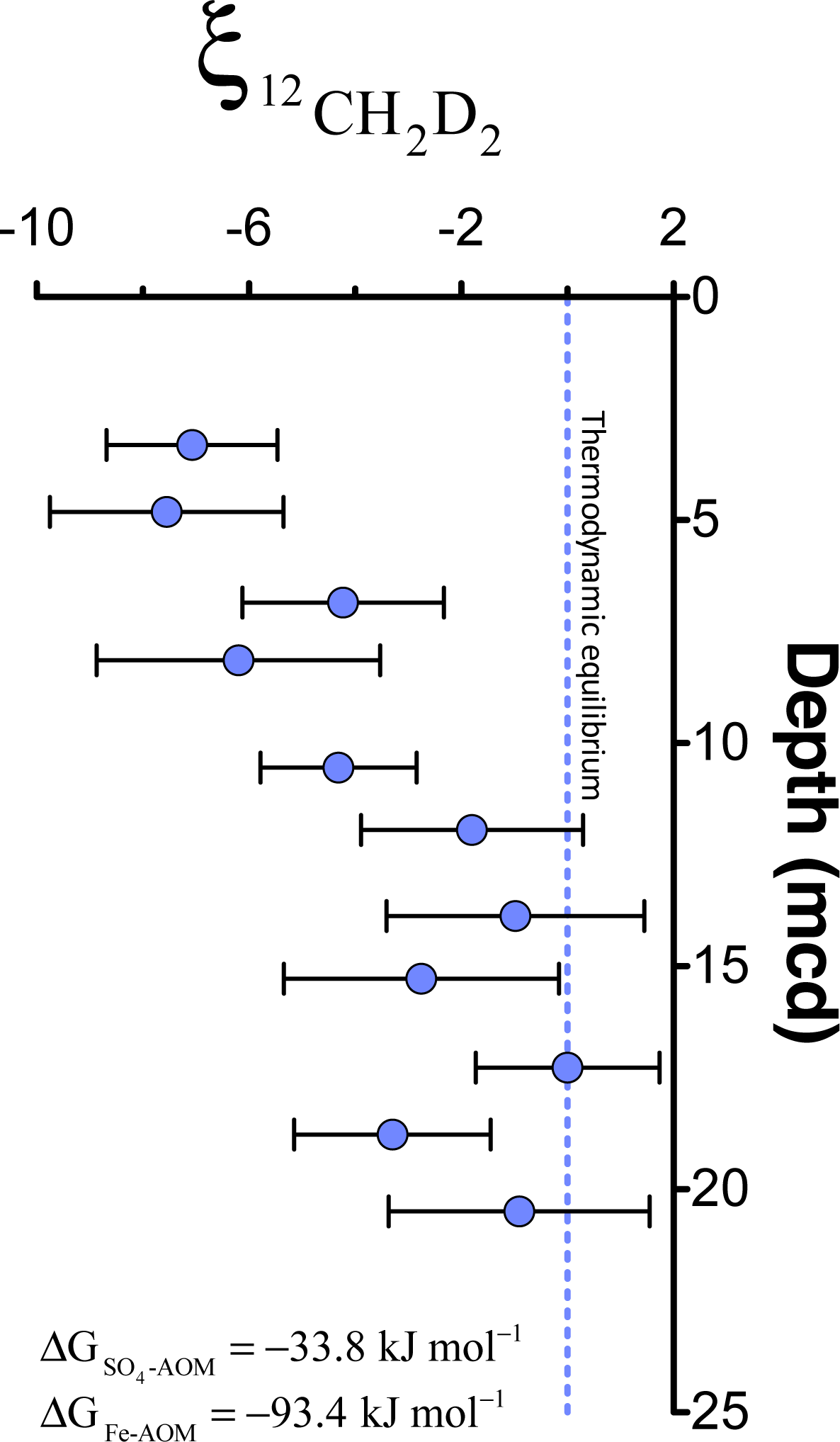
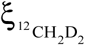
 values (±2σ) are the most negative at shallow MCD, indicating the greatest departure from equilibrium and approach zero (violet dashed line) with increasing MCD, indicating an approach towards intra-species thermodynamic equilibrium. ΔG values suggest either metabolism is capable of reversibility (See Supplemental Information).

Methanogenesis in intra-molecular isotopic thermodynamic equilibrium requires reversibility at the final hydrogen-addition step. Trace methane oxidation (TMO) does occur during methanogenesis, but altering the concentration of H_2_ or terminal electron acceptors necessary for oxidizing methane does not increase TMO and has never been shown to consume greater than ~3% of the CH_4_ produced (MORAN *et al.*, 2005; MORAN *et al.*, 2007). Although some of the first steps of methanogenesis may be reversible (VALENTINE *et al.*, 2004), the final transfer of the methyl group to coenzyme M and the subsequent final hydrogen addition are believed to be irreversible (GÄRTNER *et al.*, 1994; THAUER, 2011). Work on determining the biochemical pathways of Fe (III) reducing ANME has so far required genetically modifying a methanogen, *M. acetivorans* with the Mcr from an ANME-2 group organism because unmodified *M. acetivorans* are unable to be cultured on methane without the modified Mcr (SOO *et al.*, 2016; YAN *et al.*, 2018).

In contrast, AOM is known to have an enzymatic back reaction that suggests the first step of anaerobic methanotrophy is partially reversible, providing a potential mechanism for equilibrating methane isotopologues with environmental temperatures. This back reaction is sensitive to concentrations of the terminal electron acceptor 
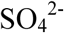
 (HOLLER *et al.*, 2011; YOSHINAGA *et al.*, 2014). At high concentrations of 
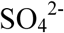
, the back reaction produces 3-7% of the CH_4_ consumed by AOM, and at low concentrations of 
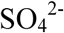
 (below 0.5 mM), back reaction can produce as much as 78% of the CH_4_ consumed by AOM (YOSHINAGA *et al.*, 2014; TIMMERS *et al.*, 2017). Intracellular exposure of methane to *Mcr* can lead to isotope exchange catalysis during bond rupture and reformation (MARLOW *et al.*, 2017), and methane that diffuses back out of the cell could move an environmental reservoir of methane towards thermodynamic equilibrium. Using estimates of AOM rates from Dijkstra et al., (2018), we calculate the timescale of this equilibration (see Supplemental Information) to be on the order of 10^3^-10^4^ years, consistent with the estimated age of methane at 20 MCD of <~8000 years [Fig. S1-S2]. Such timescales would vary in environments with differing rates of AOM.

Our findings suggest that when terminal electron acceptors are limiting, equilibrium bond ordering may be used to identify AOM activity even when the bulk isotopic composition of reservoir material is changing and ambiguous. Methane from Kidd Creek Mine (Ontario, Canada) also shows methane isotopologues transitioning with time from non-equilibrated to equilibrated (Fig. 3), suggesting that as energy-rich fractures open, AOM consumes the abiotically produced methane and drives it towards thermodynamic equilibrium (YOUNG *et al.*, 2017).

**Figure 3:**
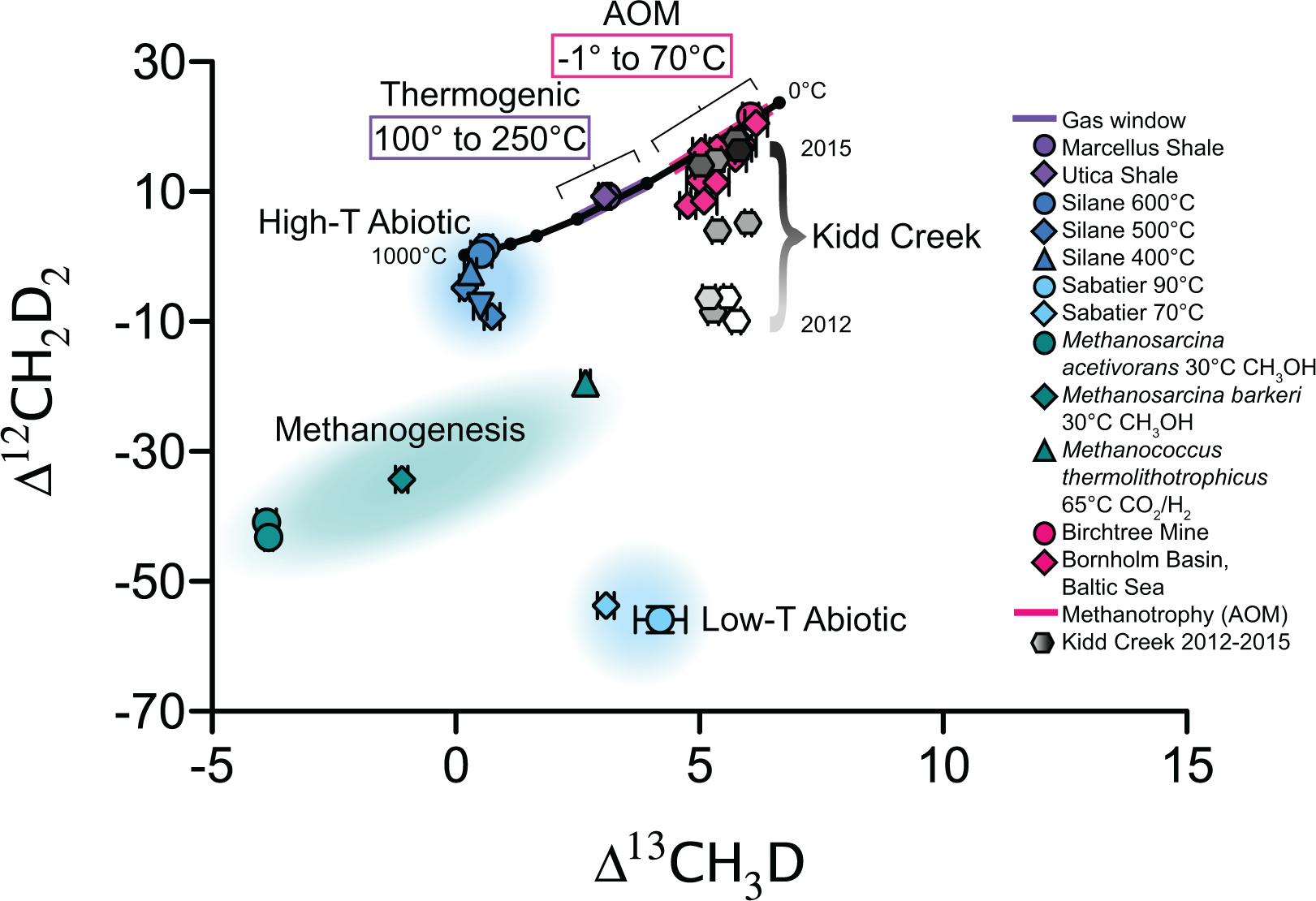
Δ^12^CH_2_D_2_ is plotted versus Δ^13^CH_3_D. The solid black line represents theoretical thermodynamic equilibrium abundances (with dots representing 100°C increments from 0-1000°C). Methane produced by thermogenesis, high-temperature abiotic reactions, microbial methanogenesis and low-temperature abiotic reactions inhabit unique zones in double-isotopologue space. AOM, equilibrating through exchange catalysis during enzymatic back reaction, also inhibits a unique zone: low-temperature intra-species thermodynamic equilibrium. Bornholm Basin data (magenta diamonds) span between the methanogenesis and AOM zones, while data from Kidd Creek Mine (black to white symbols)(YOUNG *et al.*, 2017) span from the abiotic to AOM zone. See Supplemental Information for additional sample description.

Methane driven to equilibrium by AOM has a distinct Δ^13^CH_3_D and Δ^12^CH_2_D_2_ composition when compared to methane produced through other known mechanisms (thermogenesis, microbial methanogenesis and abiotic methanogenesis) (Fig. 3). AOM metabolisms have been shown to be active from −1°C to 70°C (NIEMANN *et al.*, 2006; HOLLER *et al.*, 2011), a range of temperatures that is unique and distinguishable from equilibrated thermogenic methane forming in the 100°C to 250°C gas window. Abiotic methane may form also form at low temperatures, yet even when it is apparently equilibrated in Δ^13^CH_3_D, it may exhibit large depletions in Δ^12^CH_2_D_2_ that distinguish it from methane that has undergone exchange catalysis during AOM. We suggest that when combined (as in the 
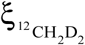
 parameter), Δ_12_CH_2_D_2_ and Δ^13^CH_3_D values are sensitive indicators of the degree of thermodynamic equilibrium that may be useful in determining the role of enzymatic back-flux during AOM in the global methane cycle.

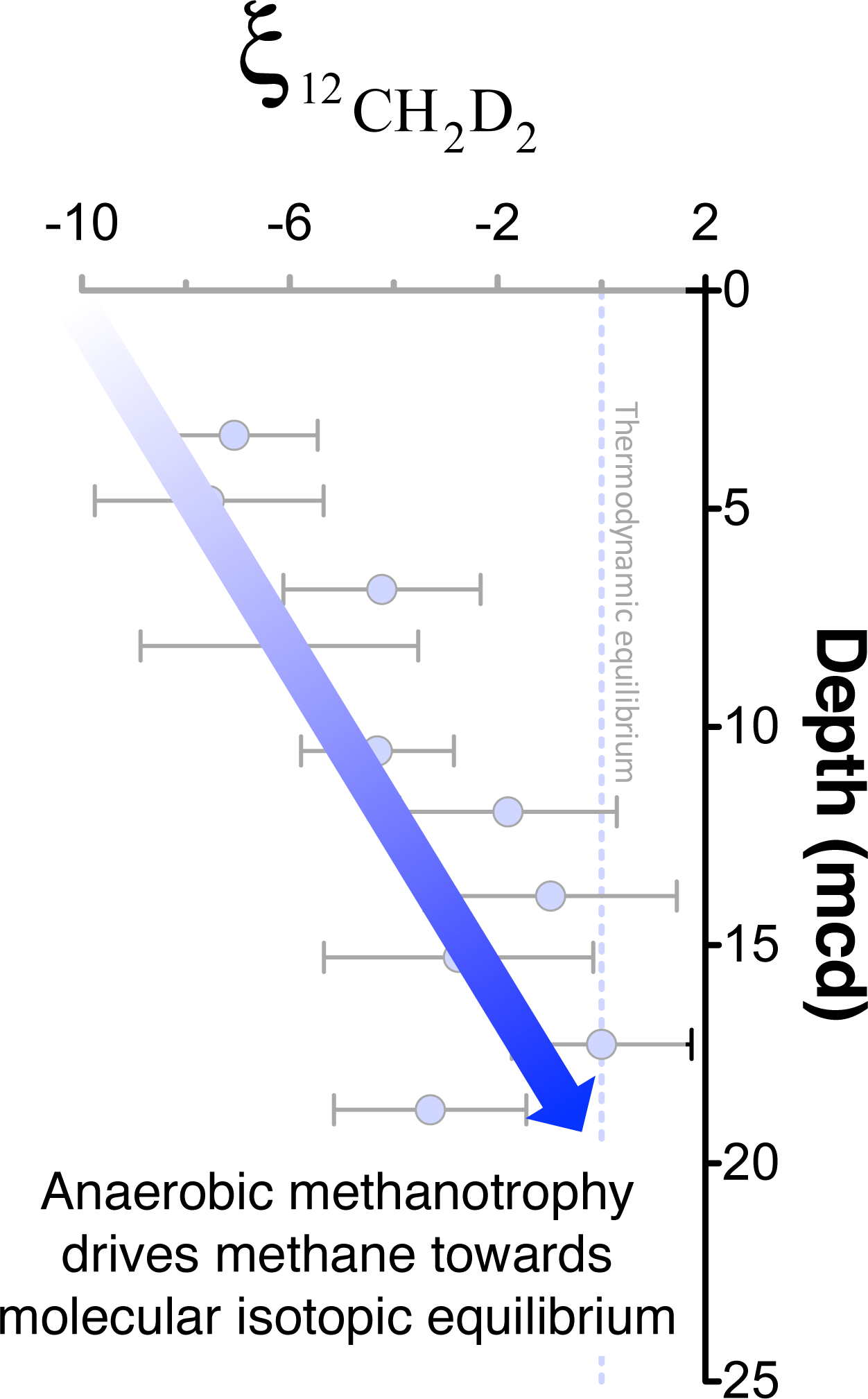

